# Effects of Nicotine and THC Vapor Inhalation Administered by An Electronic Nicotine Delivery System (ENDS) in Male Rats

**DOI:** 10.1101/418699

**Authors:** Mehrak Javadi-Paydar, Tony M. Kerr, Eric L. Harvey, Maury Cole, Michael A. Taffe

**Affiliations:** Department of Neuroscience; The Scripps Research Institute; La Jolla, CA, USA; La Jolla Alcohol Research, Inc; La Jolla, CA, USA

**Author notes:** Address Correspondence to: Dr. Michael A. Taffe, Department of Neuroscience, SP30-2400; 10550 North Torrey Pines Road; The Scripps Research Institute, La Jolla, CA 92037; USA.

## Abstract

**Background:** Electronic nicotine delivery systems (ENDS, e-cigarettes) are increasingly used for the self-administration of nicotine by various human populations, including previously nonsmoking adolescents. Studies in preclinical models are necessary to evaluate health impacts of ENDS including the development of nicotine addiction, effects of ENDS vehicles, flavorants and co-administered psychoactive substances such as ∆^9^-tetrahydrocannabinol (THC). This study was conducted to validate a rat model useful for the study of nicotine effects delivered by inhalation of vapor created by ENDS.

**Methods:** Male Sprague-Dawley rats (N=8) were prepared with radiotelemetry devices for the reporting of temperature and activity. Experiments subjected rats to inhalation of vapor generated by an electronic nicotine delivery system (ENDS) adapted for rodents. Inhalation conditions included vapor generated by the propylene glycol (PG) vehicle, Nicotine (1, 10, 30 mg/mL in the PG) and THC (12.5, 25 mg/mL).

**Results:** Nicotine inhalation increased spontaneous locomotion and decreased body temperature of rats. Pretreatment with the nicotinic cholinergic receptor antagonist mecamylamine (2 mg/kg, i.p.) prevented stimulant effects of nicotine vapor inhalation and attenuated the hypothermic response. Combined inhalation of nicotine and THC resulted in apparently independent effects which were either additive (hypothermia) or opposed (activity).

**Conclusions:** These studies provide evidence that ENDS delivery of nicotine via inhalation results in nicotine-typical effects on spontaneous locomotion and thermoregulation in male rats. Effects were blocked by a nicotinic antagonist, demonstrating mechanistic specificity. This system will therefore support additional studies of the contribution of atomizer/wick design, vehicle constituents and/or flavorants to the effects of nicotine administered by ENDS.

## Introduction

Exponential growth in the use of electronic nicotine delivery systems (ENDS or e-cigarette) for the self-administration of nicotine has been observed across numerous populations within the US (McCarthy 2013; Wong and Fan 2018) and worldwide (Brown et al. 2014; Fraser et al. 2018; Goniewicz and Zielinska-Danch 2012; Sarfraz et al. 2018; Wang et al. 2018; Yoong et al. 2018) in recent years. Evidence suggests that ENDS are being used in cessation attempts (Kalkhoran and Glantz 2016; Mantey et al. 2017; Weaver et al. 2018) as well as by people who choose to continue their nicotine habit either for potential / perceived health benefits (Spears et al. 2018; Stokes et al. 2018) or social acceptability (Berg et al. 2014; Romijnders et al. 2018) compared with tobacco smoking. Widespread availability of such devices, particularly with adolescent age ranges, sparks concern that ENDS may facilitate the development of nicotine dependence without any prior use of combustible tobacco products (Fulton et al. 2018; Martinasek et al. 2018) and may even increase the odds of later initiation of combustible tobacco use (East et al. 2018; Loukas et al. 2018; Primack et al. 2018), a relationship that may be mediated by the nicotine content in the e-cigarettes (Goldenson et al. 2017).

There are as yet relatively few studies of the effects of ENDS in rodent models, however recent studies showed e-cigarette based vapor inhalation of nicotine in mice reduced body temperature and locomotor activity (Lefever et al. 2017) and increased platelet activity (Qasim et al. 2018), which together demonstrate the feasibility of the approach. The paucity of models is underlined by the fact that several recent studies have resorted to parenteral injection of e-cigarette refill liquids to determine effects in rats (Bunney et al. 2018; El Golli et al. 2016; Golli et al. 2016; Harris et al. 2015; LeSage et al. 2016). Inhalation administration of nicotine is possible, since one study showed that inhalation of air bubbled through an aqueous solution of nicotine produces nicotine dependence and facilitates the acquisition of subsequent nicotine intravenous self-administration (Gilpin et al. 2014). In addition, a few studies have examined the effects of inhalation of vapor generated by e-cigarette devices on toxicological markers (Phillips et al. 2017; Werley et al. 2016), wound healing (Rau et al. 2017) and laryngeal mucosa (Salturk et al. 2015) in rats. This relative paucity of information means that it is of pressing and significant interest to develop and validate experimental methods that can support a range of ENDS studies. We have recently developed a system capable of delivering behaviorally significant doses of THC (Javadi-Paydar et al. 2018a; Nguyen et al. 2016b; Nguyen et al. 2018), as well as methamphetamine, mephedrone and 3,4-methylenedioxypyrovalerone (Nguyen et al. 2016a; Nguyen et al. 2017) to rats via the inhalation of ENDS vapor. This system has also been shown to support the self-administration of sufentanil with rats reaching dependence-inducing levels of drug intake via inhalation (Vendruscolo et al. 2018).

The present study was designed to first determine if the inhalation of nicotine using this vapor inhalation system produces either hypothermic or locomotor effects in rats. Injected nicotine has previously been shown to reduce body temperature and increase spontaneous locomotion in rats (Bryson et al. 1981; Clemens et al. 2009; Green et al. 2003; Levin et al. 2003), thus these measures were selected for initial validation. These have the additional advantages of supporting indirect cross-drug comparison with our prior reports on the effects of inhaled THC (Javadi-Paydar et al. 2018a; Nguyen et al. 2016b; Nguyen et al. 2018) and psychomotor stimulants (Nguyen et al. 2016a; Nguyen et al. 2017). As in those prior studies, it was further important to contextualize some of the inhaled effects, including plasma nicotine and cotinine levels, with those produced by parenteral nicotine injection.

A second goal was to determine if there were interactive effects of the inhalation of threshold doses of THC and nicotine. Many consumers use hollowed out cigars filled with cannabis (“blunts”) to dose themselves with THC (Eggers et al. 2017; Ream et al. 2008; Schauer et al. 2017; Timberlake 2009) and other individuals co-use tobacco and marijuana sequentially in a manner that would expose them simultaneously to both nicotine and THC (Agrawal et al. 2012). This practice may leave some users at higher risk for cannabis dependence (Ream et al. 2008). The burgeoning use of e-cigarette devices to consume THC via cannabis extracts makes it trivial for users to titrate drug combinations if desired. It can be difficult to identify independent versus additive risks from heterogenous human populations who use both tobacco and cannabis; well controlled preclinical models can be useful for identifying independent or interactive effects of nicotine and THC co-administration. Therefore additional studies were designed to determine if vapor inhalation of THC modified the effects of nicotine inhalation.

## Methods

### Subjects

Male Sprague-Dawley (N=8; Harlan/Envigo, Livermore, CA) rats were housed in humidity and temperature-controlled (23±2 °C) vivaria on 12:12 hour light:dark cycles. Rats had *ad libitum* access to food and water in their home cages and all experiments were performed in the rats’ scotophase. Rats entered the laboratory at 10-11 weeks of age. All procedures were conducted under protocols approved by the Institutional Animal Care and Use Committee of The Scripps Research Institute.

### Drugs

Rats were exposed to vapor derived from nicotine bitartrate (1, 10, 30 mg/mL) or ∆9-tetrahydrocannabinol (THC; 12.5, 25 mg/mL) dissolved in a propylene glycol (PG) vehicle. Doses for nicotine were derived from retail products intended for human use and interval reductions found to produce differential effects in pilot studies in the laboratory. THC concentrations were likewise selected from our prior/ongoing work, in this case to fit the intent to achieve threshold doses for evaluation of potential drug interactions. The ethanolic THC stock was aliquoted in the appropriate volume, the ethanol evaporated off and the THC was then dissolved in the PG to achieve target concentrations for vaporized delivery. Four 10-s vapor puffs were delivered with 2-s intervals every 5 minutes, which resulted in use of approximately 0.1 mL in a 30 minutes exposure session. THC (5 mg/kg) or nicotine bitartrate (0.4 mg/kg) were injected, i.p., in a 1:1:8 ratio of ethanol:cremulphor:saline. Mecamylamine HCl (2 mg/kg) and nicotine bitartrate (0.8, 2.0 mg/kg) were dissolved in saline for subcutaneous injection, selection of these doses was from relevant precedent literature. Nicotine and mecamylamine were obtained from Sigma-Aldrich (St. Louis, MO) and the THC was provided by the U.S. National Institute on Drug Abuse.

### Radiotelemetry

Sprague-Dawley rats (N=8) were anesthetized with an isoflurane/oxygen vapor mixture (isoflurane 5% induction, 1-3% maintenance) and sterile radiotelemetry transmitters (Data Sciences International, St. Paul, MN; TA-F40) were implanted in the abdominal cavity through an incision along the abdominal midline posterior to the xyphoid space as previously described (Javadi-Paydar et al. 2018b). Experiments were initiated four weeks after surgery. Activity and temperature responses were evaluated in the vapor inhalation chambers in a dark testing room, separate from the vivarium, during the (vivarium) dark cycle. The telemetry recording plates were placed under the vapor inhalation chambers thus data were collected on a 5 minute schedule throughout the experiment.

### Inhalation Apparatus

Sealed exposure chambers were modified from the 259mm X 234mm X 209mm Allentown, Inc (Allentown, NJ) rat cage to regulate airflow and the delivery of vaporized drug to rats using e-cigarette cartridges (Protank 3 Atomizer, MT32 coil operating at 2.2 ohms, by Kanger Tech; Shenzhen Kanger Technology Co.,LTD; Fuyong Town, Shenzhen, China) as has been previously described (Nguyen et al. 2016a; Nguyen et al. 2016b). An e-vape controller (Model SSV-1; 3.3 volts; La Jolla Alcohol Research, Inc, La Jolla, CA, USA) was triggered to deliver the scheduled series of puffs by a computerized controller designed by the equipment manufacturer (Control Cube 1; La Jolla Alcohol Research, Inc, La Jolla, CA, USA). The chamber air was vacuum controlled by a chamber exhaust valve (i.e., a “pull” system) to flow room ambient air through an intake valve at ~1 L per minute. This also functioned to ensure that vapor entered the chamber on each device triggering event. The vapor stream was integrated with the ambient air stream once triggered.

### Plasma Nicotine and Cotinine Analysis

Plasma nicotine and cotinine content was quantified using liquid chromatography/mass spectrometry (LCMS). Briefly, 50 ul of plasma were mixed with 50 ul of deuterated internal standard (100 ng/ml cotinine-d3 and nicotine-d4; Cerilliant). Nicotine and cotinine (and the internal standards) were extracted into 900 ul of acetonitrile and then dried. Samples were reconstituted in 100 uL of an acetonitrile/water (9:1) mixture. Separation was performed on an Agilent LC1200 with an Agilent Poroshell 120 HILIC column (2.1mm x 100mm; 2.7 um) using an isocratic mobile phase composition of acetonitrile/water (90:10) with 0.2% formic acid at a flow rate of 325 uL/min. Nicotine and cotinine were quantified using an Agilent MSD6130 single quadrupole interfaced with electrospray ionization and selected ion monitoring [nicotine (m/z=163.1), nicotine-d4 (m/z=167.1), cotinine (m/z=177.1) and cotinine-d3 (m/z=180.1)]. Calibration curves were generated daily using a concentration range of 0-200 ng/mL with observed correlation coefficients of 0.999.

### Experiments

#### Experiment 1 (Nicotine Dose)

The effects of inhaling three concentrations of nicotine (1, 10, 30 mg/mL) were compared with the inhalation of the PG vehicle. A single 30 minute inhalation interval, selected based on pilot studies, was conducted each test session and doses were evaluated in a balanced order with 3-4 days minimum between test days for this and all subsequent experiments. For comparison, the effects of injected nicotine (0, 0.8 mg/kg, s.c.) were assessed with the dose conditions administered in a counter-balanced order. This latter experiment was conducted after Experiment 4.

#### Experiment 2 (Repeated Nicotine Inhalation)

Since human tobacco smokers tend to smoke multiple times per day, an experiment was conducted to determine the effects of inhaling PG vs nicotine (30 mg/mL) for 15 minutes every hour for four total inhalation bouts. Following Experiment 4, this condition was repeated, including pretreatments of saline versus mecamylamine (2 mg/kg, i.p.) for four total conditions, assessed in a counter-balanced order. In this latter experiment pretreatments were administered 15 minutes prior to the start of inhalation, i.e., in the middle of the usual baseline interval.

#### Experiment 3 (Nicotine combined with THC)

The effects of inhaling vapor from PG, nicotine (30 mg/mL), THC (25 mg/mL) or the combination of drugs in a 30 minute session were next evaluated, in a counter-balanced order. For comparison, the effects of injected nicotine (0, 0.4 mg/kg, i.p.) in combination with injected THC (0, 5 mg/kg, i.p.) were assessed with the dose conditions administered in a counter-balanced order (nicotine was injected i.p. in this experiment to avoid multiple injections for the combination condition). This latter experiment was conducted after Experiment 5.

#### Experiment 4 (Repeated nicotine combined with THC)

The effects of inhaling vapor from PG, nicotine (30 mg/mL), THC (12.5 mg/mL) or the combination of drugs for 15 minutes every hour, for four total inhalation epochs, was assessed in this experiment. The inhalation / dose conditions were administered in a counter-balanced order.

#### Experiment 5 (Plasma nicotine and cotinine after vapor inhalation)

Blood samples (~0.5 ml) were withdrawn from the jugular vein under inhalation anesthesia following sessions of vapor inhalation of nicotine (30 mg/mL) or injection of nicotine (0.2, 0.4, 0.8 mg/kg, s.c.). Animals in the telemetry group were assessed once after a 15 minute session and once after a 30 minute session in a balanced order.

### Data Analysis

Temperature and activity rate (counts per minute) were collected via the radiotelemetry system on a 5-minute schedule and analyzed in 30 minute averages (the time point refers to the ending time, i.e. 60 = average of 35-60 minute samples). In the mecamylamine pre-treatment study, the baseline interval is depicted as two 15-minute bins with the s.c. injection just prior to the second baseline interval. Any missing temperature values were interpolated from surrounding values, this amounted to less than 10% of data points. Telemetry data were analyzed with Analysis of Variance (ANOVA) with repeated measures factors for the Drug Condition and the Time post-initiation of vapor or Time post-injection. Comparison with the pre-treatment baselines and the vehicle conditions Plasma levels of nicotine and cotinine were analyzed with ANOVA with repeated measures factors of Drug Condition and Time post-initiation of vapor or Time post-injection. A between-subjects factor was included for the strain comparison. Any significant effects within group were followed with post-hoc analysis using Tukey correction for all multi-level, and Sidak correction for any two-level, comparisons. All analysis used Prism 7 for Windows (v. 7.03; GraphPad Software, Inc, San Diego CA).

## Results

### Experiment 1 (Nicotine Dose)

The inhalation of nicotine in a 30 minute exposure reduced the body temperature of rats as is depicted in **Figure 1A**. Statistical analysis of the body temperature confirmed significant effects of Time [F (7, 49) = 49.93; P<0.0001], Drug Condition [F (3, 21) = 5.41; P<0.01] and the interaction of factors [F (21, 147) = 7.75; P<0.0001]. The Tukey post-hoc test further confirmed that body temperature was significantly lower than the pre-treatment baseline and the respective time during the PG condition only in the nicotine (30 mg/mL) condition (30-120 minutes after the initiation of vapor). In this study, locomotor activity declined across the session (significant main effect of Time F (7, 49) = 56.59; P<0.0001) to a similar extent in all treatment conditions (**Figure 1B**). The injected nicotine study (**Figure 1C,D**) confirmed that 0.8 mg/kg nicotine, s.c., reduces body temperature (Time: F (7, 49) = 33.13; P<0.0001; Drug Condition: F (1, 7) = 44.21; P<0.0005; Interaction: F (7, 49) = 19.58; P<0.0001) and activity rate (Time: F (7, 49) = 8.51; P<0.0001; Drug Condition: n.s.; Interaction: F (7, 49) = 3.56; P<0.005).

**Figure 1:**
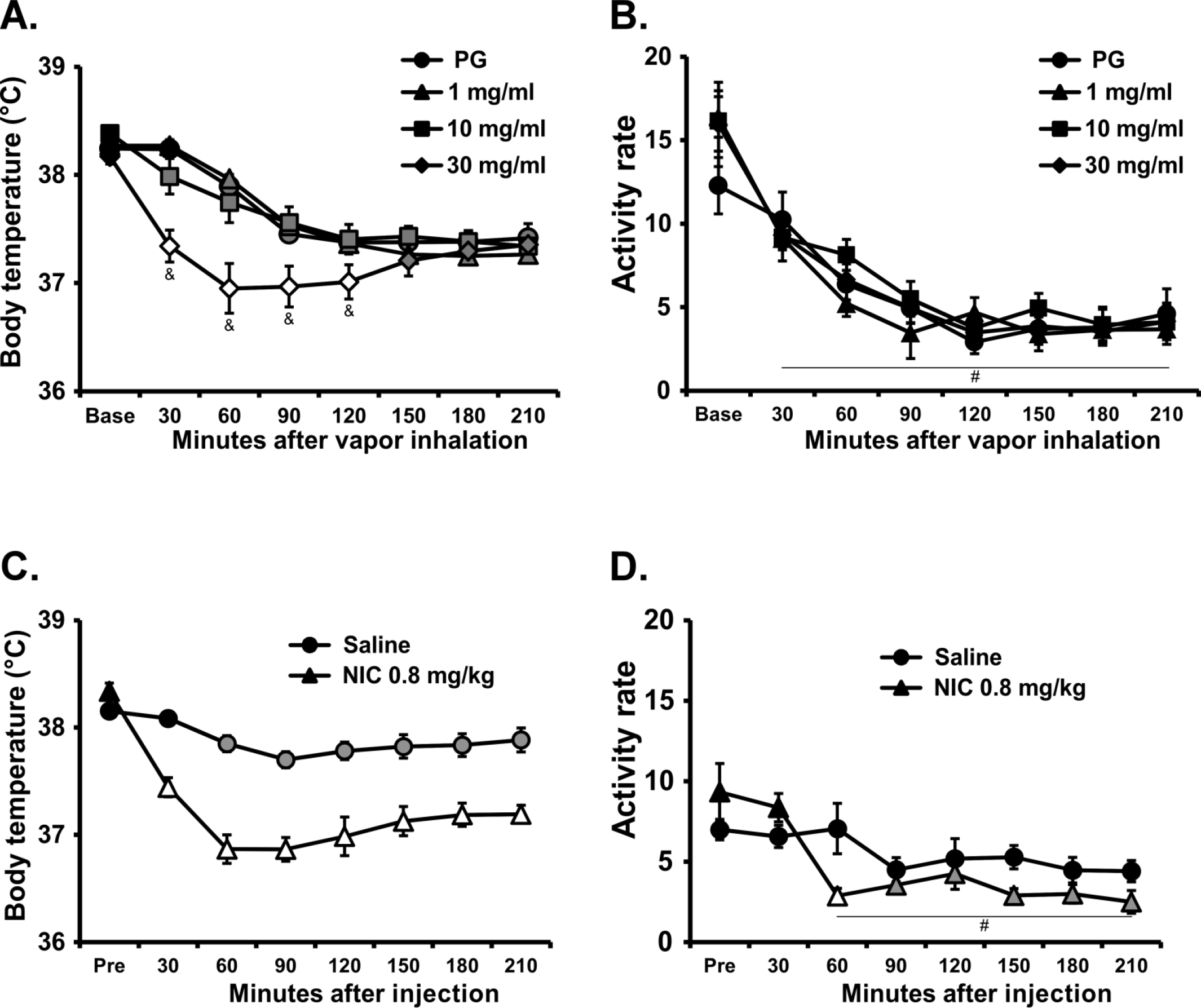
Mean (N=8; ±SEM) A,C) body temperature and B,D) activity rates after A,B) inhalation of the PG vehicle or nicotine (NIC; 1, 10, 30 mg/mL) or C,D) subcutaneous injection of saline or 0.8 mg/kg nicotine. Open symbols indicate a significant difference from both the vehicles at a given time-point and the within-treatment baseline, while shaded symbols indicate a significant difference from the baseline, within treatment condition, only. A significant difference from both 1 and 10 mg/mL NIC inhalation conditions is indicated by &. A significant difference from the baseline, collapsed across treatment condition is indicated with #.

### Experiment 2 (Repeated Nicotine Inhalation)

The inhalation of nicotine (30 mg/mL) in four 15-minute sessions separated by an hour reduced rat’s body temperature as is depicted in Figure 2A. Statistical analysis confirmed significant effects of Time [F (9, 63) = 16.7; P<0.0001] and the interaction of Drug Condition with Time [F (9, 63) = 2.83; P<0.01] on body temperature. The post-hoc test confirmed significant nicotine related hypothermia was observed from 180-240 minutes after the start of the first vapor inhalation session. Locomotor activity was likewise significantly altered by repeated nicotine inhalation (**Figure 2B**). The ANOVA confirmed significant effects of Time [F (9, 63) = 22.44; P<0.0001] and Drug Condition [F (1, 7) = 16.67; P<0.005], and of the interaction of factors [F (9, 63) = 2.55; P<0.05] on locomotor activity. The post-hoc test further confirmed that significant increases in activity relative to the pre-vapor baseline were observed after each of the nicotine inhalation intervals and relative to the vehicle condition for the second, third and fourth nicotine inhalations.

**Figure 2:**
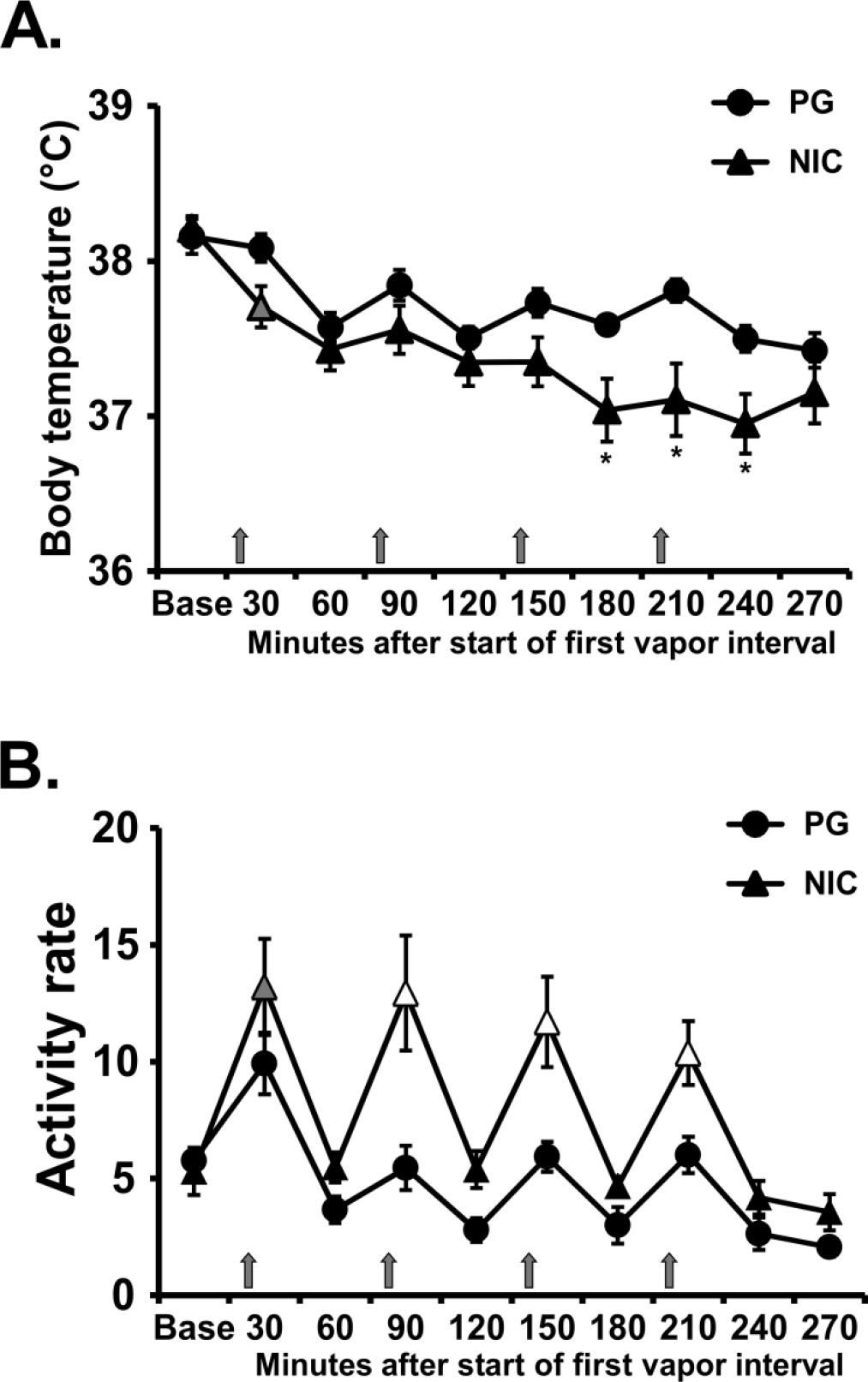
Mean (N=8; ±SEM) A) temperature and B) locomotor activity after inhalation of vapor from the propylene glycol (PG) vehicle or nicotine (NIC; 30 mg/mL) in four successive 15 minute intervals (indicated with arrows). Grey symbols indicate a significant difference from the point time point immediately prior to each inhalation interval within condition. Significant differences from the PG condition (only) are indicated with *. Open symbols indicate significant difference from preceding timepoint and PG. Base = baseline.

The administration of mecamylamine (2 mg/kg, i.p.) 15 minutes prior to the start of inhalation modified the effects of nicotine (**Figure 3**). The ANOVA confirmed significant effects on temperature (Time: F (10, 70) = 15.65; P<0.0001; Drug Condition: F (3, 21) = 5.35; P=0.0068; Interaction of factors: F (30, 210) = 3.53; P<0.0001) and on activity (Time: F (10, 70) = 13.91; P<0.0001; Drug Condition: F (3, 21) = 9.27; P<0.0005; Interaction of factors: F (30, 210) = 2.67; P<0.0001). The post-hoc test confirmed the locomotor response to nicotine epochs was prevented by mecamylamine pre-treatment (**Figure 3B**) and a significant attenuation of the hypothermic response to nicotine was confirmed after the fourth vapor interval (**Figure 3A**).

**Figure 3:**
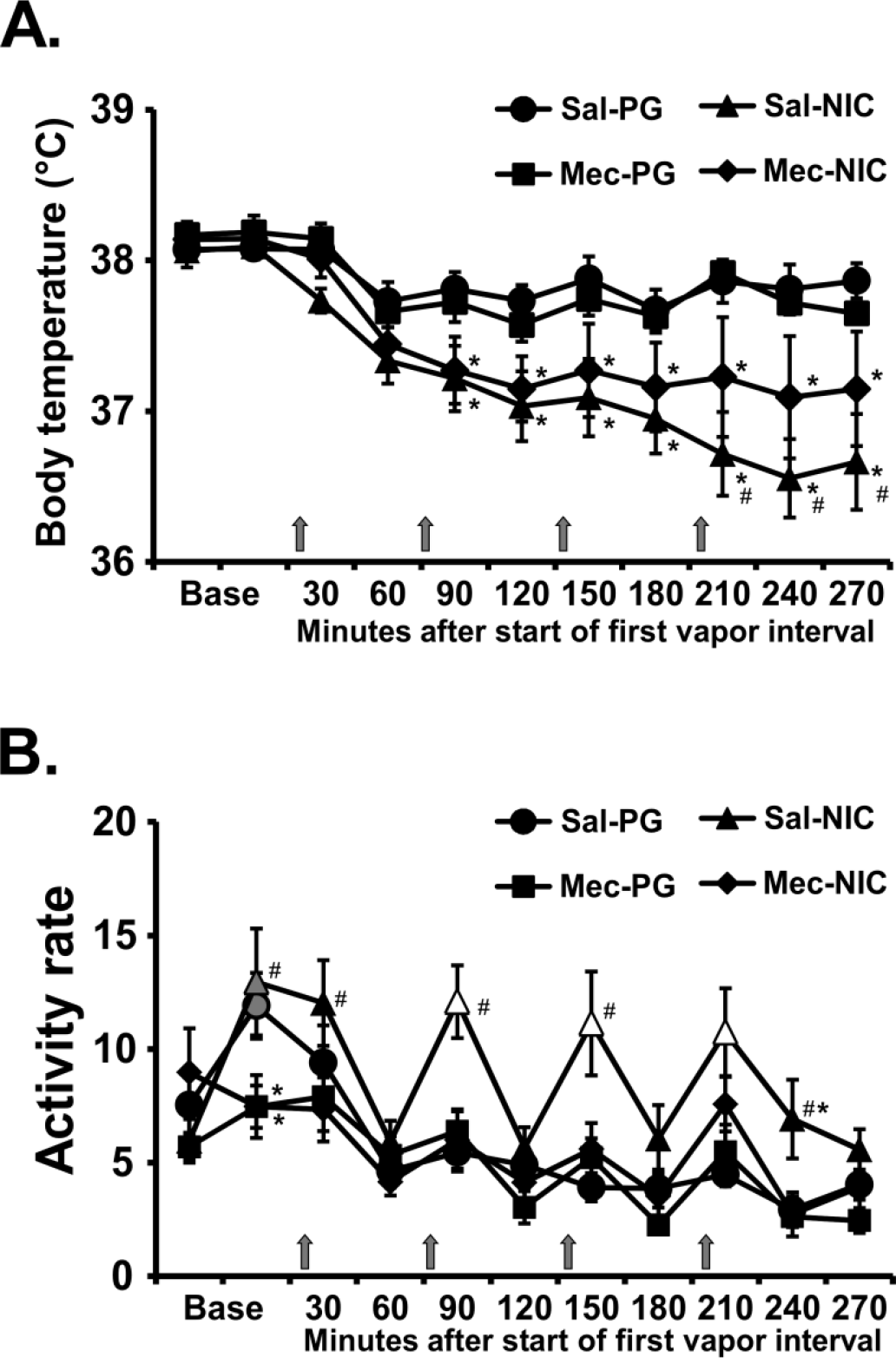
Mean (N=8; ±SEM) A) temperature and B) locomotor activity after inhalation of vapor from the propylene glycol (PG) vehicle or nicotine (NIC; 30 mg/mL) in four successive 15 minute intervals (indicated with arrows) with either mecamylamine (Mec) or Saline (Sal) i.p. 15 min prior to the first inhalation interval. Open symbols indicate a significant difference from both PG at the same time point and the point time point immediately prior to each inhalation interval within condition. Significant differences from the PG condition (only) are indicated with *. Significant differences between Saline and Mecamylamine pre-treatment within a vapor inhalation condition are indicated with #. Base = baseline (divided into two 15 minute timepoints).

### Experiment 3 (Nicotine combined with THC)

The inhalation of nicotine (30 mg/mL), THC (25 mg/mL) or the combination in a 30 minute session reduced the rats’ body temperature, as is depicted in **Figure 4A**. The statistical analysis confirmed significant effects of Time [F (7, 49) = 28.3; P<0.0001], Drug Condition [F (3, 21) = 8.50; P<0.001] and the interaction of factors [F (21, 147) = 6.79; P<0.0001] on body temperature. The Tukey post-hoc test confirmed that body temperature was significant lower than the pre-treatment baseline and the respective time during the PG condition for all active dose conditions. Furthermore there was a significant difference in temperature in the combined condition compared with THC or nicotine administered alone.

**Figure 4:**
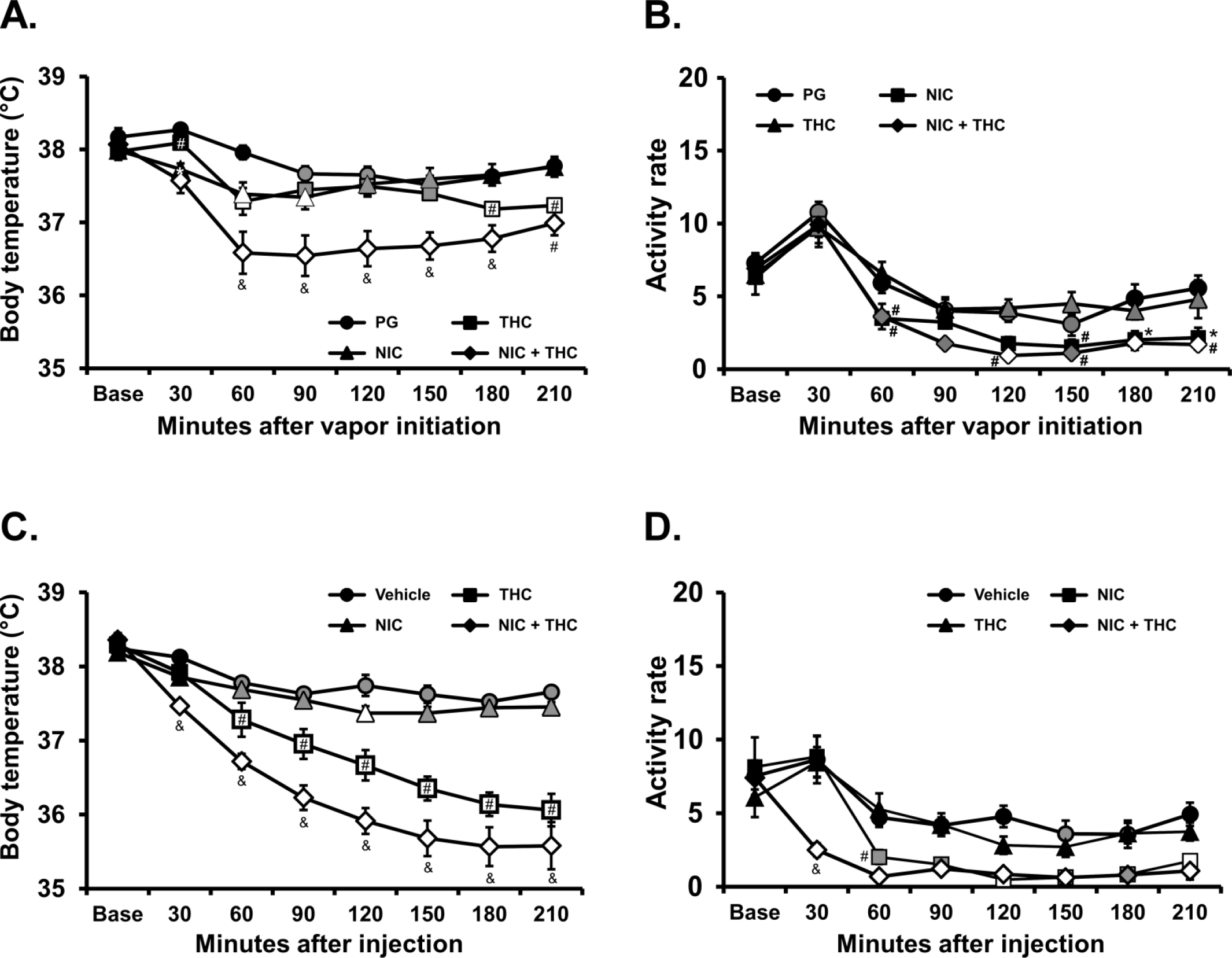
Mean (N=8; ±SEM) A) temperature and B) activity rate after inhalation of vapor from the PG vehicle, nicotine (NIC; 30 mg/mL), ∆^9^-tetrahydrocannabinol (THC; 25 mg/mL) or the combination for 30 minutes. Mean (N=8; ±SEM) C) temperature and D) activity rate after injection of nicotine (NIC; 0.4 mg/kg, i.p.), ∆^9^-tetrahydrocannabinol (THC; 5 mg/kg, i.p.) or the combination. Open symbols indicate a significant difference from both vehicle at a given time-point and the within-treatment baseline, while shaded symbols indicate a significant difference from the baseline only. Significant differences from the PG condition (only) are indicated with *. A significant difference from both NIC and THC conditions is indicated by & and from NIC by #.

The statistical analysis also confirmed significant effects of Time [F (7, 49) = 55.82; P<0.0001] and of Drug Condition [F (3, 21) = 14.63; P<0.0001] on activity rate (**Figure 4B**). The post hoc test confirmed significant reductions in activity relative to the pre-vapor baseline and the PG condition at the same time point after nicotine + THC (120, 180 and 210 minutes after the start of vapor). Activity was also lower compared with the same time point following PG inhalation after THC inhalation alone (180-210 minutes after vapor initiation).

There was a similar outcome in the injection study. The analysis confirmed first that there were significant effects of Time [F (7, 49) = 118.8; P<0.0001], Drug Condition [F (3, 21) = 37.95; P<0.0001] and the interaction of factors [F (21, 147) = 16.86; P<0.0001] on body temperature (**Figure 4C**). The post-hoc test confirmed that body temperature was significant lower than the pre-treatment baseline and the respective time during the PG condition for all active dose conditions, although the effect of nicotine was limited to 210 minutes after injection. The temperature in the combined treatment differed significantly from all other treatments and temperature after THC injection differed significantly from temperature after nicotine injection. The analysis also confirmed that there were significant effects of Time [F (7, 49) = 36.12; P<0.0001], Drug Condition [F (3, 21) = 14.39; P<0.0001] and the interaction of factors [F (21, 147) = 2.57; P<0.001] on activity rate (**Figure 4D**). The post-hoc test confirmed that activity rate was significantly suppressed compared with the baseline in the THC or THC + nicotine injection conditions and suppressed relative to the vehicle injection condition after the combination.

### Experiment 4 (Repeated Nicotine combined with THC)

Repetition of four 15 minute inhalation intervals resulted in detectable effects of nicotine, THC and the combination of the two (**Figure 5A**). The analysis confirmed significant effects of Time [F (9, 63) = 32.53; P<0.0001], Drug Condition [F (3, 21) = 4.63; P<0.05] and the interaction of factors [F (27, 189) = 2.63; P<0.0001] on body temperature. The post hoc test further confirmed that body temperature was lower during the combined condition compared with the PG or nicotine inhalation 90-270 minutes after the start of the first inhalation period. Temperature was also reduced in the THC (12.5 mg/mL) condition, compared with PG or nicotine inhalation from 180-270 minutes after the start of the first inhalation. Spontaneous locomotor activity was increased by both THC and nicotine inhalation (**Figure 5B**). The analysis confirmed that activity rate was significantly affected by Time [F (9, 63) = 36.89; P<0.0001], Drug Condition [F (3, 21) = 10.56; P<0.0005] and the interaction of factors [F (27, 189) = 2.86; P<0.0001]. The post hoc test confirmed that activity was elevated relative to the pre-vapor time point and corresponding PG time point after the first THC exposure, the second NIC+THC exposure and the third and fourth nicotine exposures.

**Figure 5:**
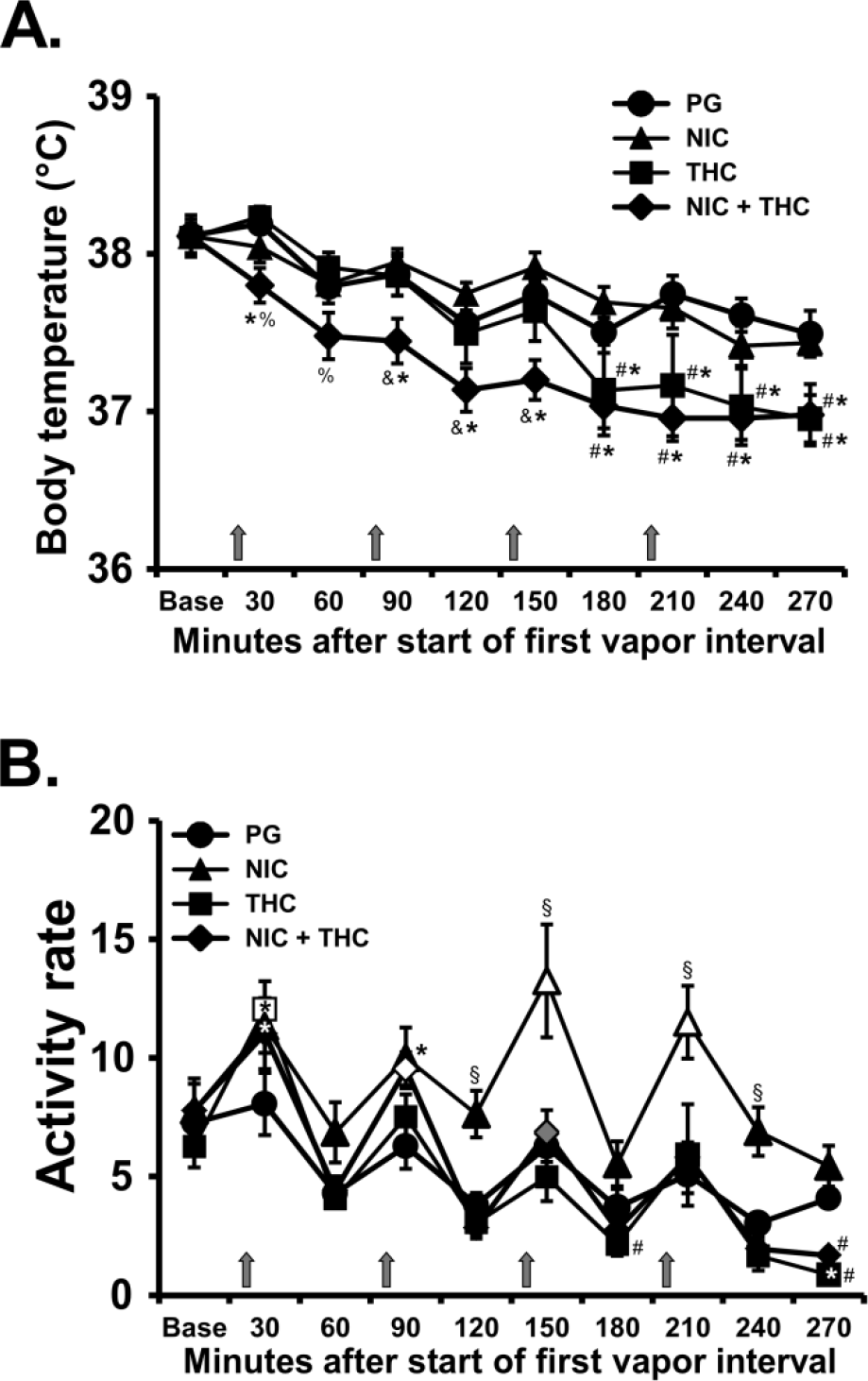
Mean (N=8; ±SEM) A) temperature and B) activity after inhalation of vapor from the PG vehicle, nicotine (NIC; 30 mg/mL), ∆9-tetrahydrocannabinol (THC; 12.5 mg/mL) or the combination, in four successive 15 minute intervals (indicated with arrows). Open symbols indicate a significant difference from both PG at the same time point and the point time point immediately prior to each inhalation interval within condition. A significant difference from the preceding time point within condition (only) is indicated with gray symbols. A significant difference from both NIC and THC conditions at the respective time point is indicated by &, from THC and the NIC+THC combination by §, from PG by *, from THC by % and from NIC by #.

### Experiment 5 (Plasma nicotine and cotinine after vapor inhalation)

Inhalation of nicotine vapor produced inhalation-duration-dependent effects on plasma nicotine and cotinine in the rats (**Figure 6A**). The ANOVA confirmed significant effects of inhalation Duration [F (1, 7) = 28.48; P<0.005] and analyte [F (1, 7) = 5.79; P<0.05] but not of the interaction of factors. The follow up study (**Figure 6B**) confirmed that plasma nicotine was higher than plasma cotinine 15 minutes after injection of nicotine and higher after the 0.8 mg/kg compared with the 2.0 mg/kg dose. The ANOVA (between subjects since a sample for one subject was unavailable for the 2.0 dose) confirmed significant effects of Dose [F (1, 13) = 85.07; P<0.0001], of Analyte [F (1, 13) = 121.1; P<0.0001] and the interaction of Dose with Analyte [F (1, 13) = 59.09; P<0.0001].

**Figure 6:**
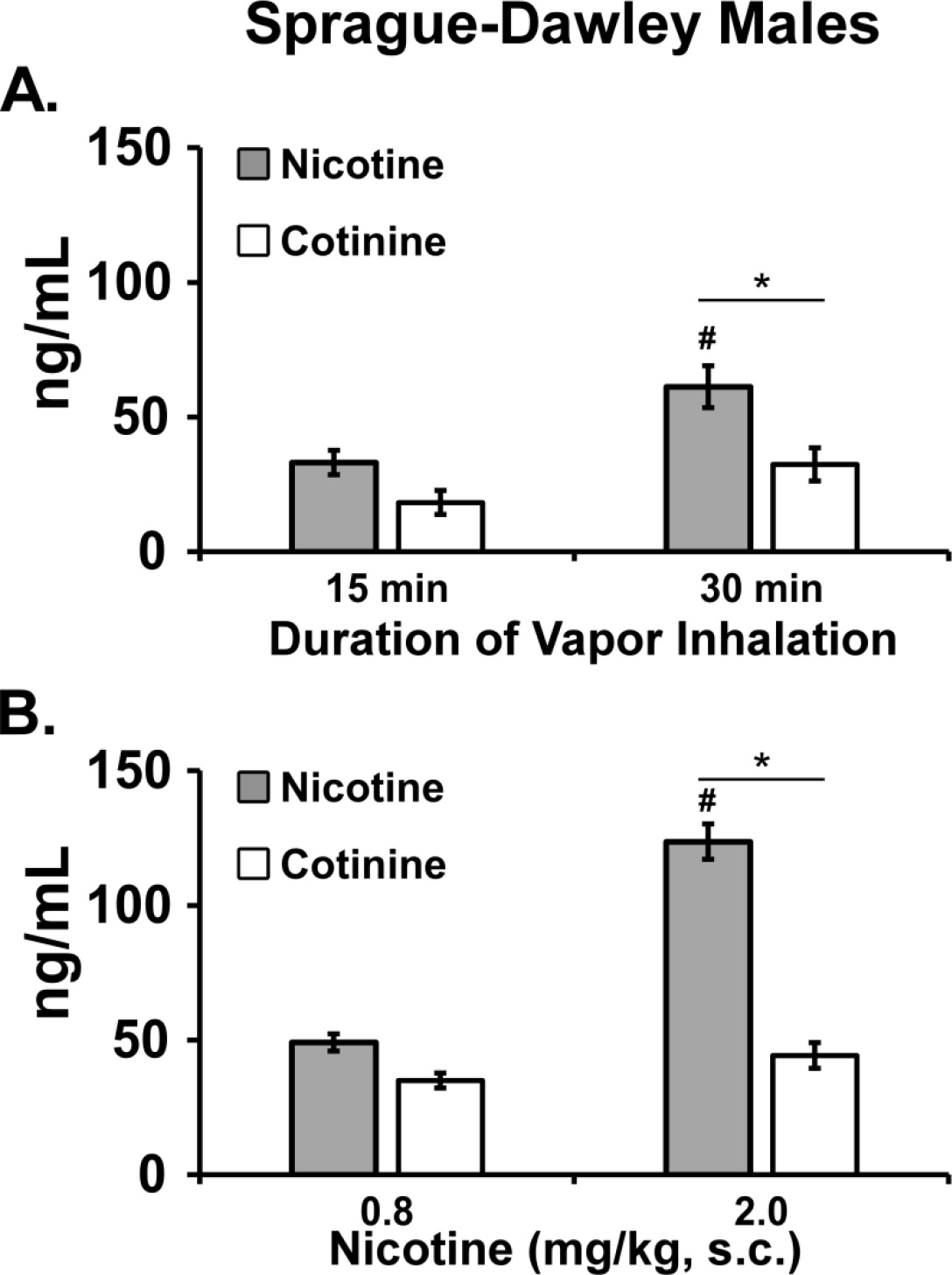
A) Mean (N=8; ±SEM) plasma nicotine and cotinine concentrations after inhalation of vapor from nicotine (30 mg/mL) after 15 and 30 minute inhalation intervals in the telemeterized S-D rat group at 35 weeks. B) Mean (N=7-8; ±SEM) plasma nicotine and cotinine concentrations in the Sprague-Dawley rats after nicotine injection (0.8, 2.0 mg/kg, s.c.). A significant difference between doses, within analyte is indicated by # and a difference between analytes, within dose, by *.

## Discussion

There has been a growing popularity and availability of electronic nicotine delivery systems (ENDS) in recent years, which reaches across adult and adolescent populations. Indeed, the latest data from the Monitoring the Future project in the United States show a doubling in the percentage of 12th grade students who vaped nicotine in the past month (Prieur 2018) in 2018 compared with 2017. This popularity has driven a current focus of the United States Food and Drug Administration on regulating ENDS devices (Benowitz and Henningfield 2018; Farber et al. 2018) but relatively few published studies validate methods for pre-clinical investigations.

We therefore designed a study to validate a recently described e-cigarette based inhalation system for evaluating nicotine vapor exposure in rats. This study found that behaviorally significant amounts of nicotine were delivered to male Sprague-Dawley rats, resulting in both hyperactivity and hypothermia responses. The initial experiments found that a 30 mg/mL concentration in the vapor vehicle was necessary to produce robust effects (**Figure 1**) and a hypothermic response was found after either a single 30 minute exposure or four 15 minute exposures administered at hourly intervals. The magnitude of temperature reduction was comparable to that produced by 0.8 mg/kg nicotine injected subcutaneously. Hyperlocomotor responses in this model were observed most consistently with a repeated 15 minute exposure every hour (**Figure 2**), compared with a single 30 minute inhalation interval. This was most likely due to an increase in locomotor activity that followed the first vehicle (propylene glycol; PG) delivery of a given session, thus obscuring locomotor effects on nicotine in the single-exposure paradigm. This response to the vehicle was attenuated on subsequent vapor deliveries within the same session in the quadruple exposure, thus the pharmacological effect of the nicotine inhalation was unmasked. Interestingly, a 0.8 mg/kg dose of nicotine injected, s.c., *reduced* activity rates. Hypothermia is produced by parenteral injection of nicotine in rats (Levin et al. 2003) as is hyperlocomotion (Bryson et al. 1981; Clemens et al. 2009; Green et al. 2003) and decreased body temperature was observed after vapor inhalation of nicotine in mice (Lefever et al. 2017). Furthermore, the effects of inhaled nicotine in this study were shown to be mediated by the nicotinic acetylcholine receptor (nAChR) since they were attenuated (temperature) or blocked entirely (locomotion) by pre-treatment with mecamylamine (**Figure 3**). Thus, these data show that physiologically significant doses of nicotine were delivered to the rats using this method.

The plasma analysis found that nicotine and cotinine levels 15 minutes after subcutaneous injection of 0.8 mg/kg were similar to those observed after 15 minutes of inhalation of vapor from 30 mg/mL in this study. The nicotine and cotinine levels were similar to intracerebral dialysate levels of nicotine and cotinine after subcutaneous injection of 0.7 mg/kg in Wistar rats (Katner et al. 2015) and plasma levels after a 0.03 mg/kg intravenous injection in Wistar rats (de Villiers et al. 2004). Concentrations of nicotine and cotinine in the plasma observed after 30 minutes of inhalation were significantly higher than those observed after 15 minutes of inhalation, confirming the control of dose via inhalation duration. Comparison of the plasma levels reached and the hypothermia responses after 0.8 mg/kg, nicotine, s.c., with those after 30 minutes of nicotine vapor (30 mg/mL; **Figure 1**) inhalation echoes a prior result in which plasma methamphetamine levels observed after inhalation and injection conditions that produced equivalent locomotor stimulation differed substantially (Nguyen et al. 2017).

Due to reasonably frequent co-exposure of human cannabis users to nicotine (Cooper and Haney 2009; Eggers et al. 2017; Ream et al. 2008; Schauer et al. 2017; Timberlake 2009), this study further determined the effects of co-exposure to THC and nicotine via vapor inhalation. Prior work with our inhalation model demonstrated consistent, dose-dependent hypothermic effects of THC inhalation and an inconsistent suppression of locomotor behavior in male and female Wistar, as well as male Sprague-Dawley, rats (Javadi-Paydar et al. 2018a; Nguyen et al. 2016b; Nguyen et al. 2018). Parenteral injection of nicotine has previously been found to increase the hypothermia associated with injected THC (Pryor et al. 1978; Valjent et al. 2002), thus there was reason to expect similar effects in this model.

Overall, the present data are most consistent with independent and additive, rather than interactive, effects of the two drugs. First, the body temperature observed after a single 30 minute exposure decreased more rapidly after nicotine (30 mg/mL) inhalation and less rapidly after THC (25 mg/mL) inhalation, but reached the same nadir. When the combination of drugs was available in the single 30 minute exposure epoch (**Figure 4**), the initial hypothermia was rapid and of the same magnitude as the response to nicotine alone in the first 30 minutes. An additional decrease over the next 30 minutes appeared to parallel the effect of THC alone, albeit starting from the lower, nicotine-induced temperature. This pattern was repeated in the injection study even though the time course of effects of THC delivered by i.p. injection is much longer than when delivered by inhalation (see (Nguyen et al. 2016b; Taffe et al. 2015)). In the study in which four 15-minute inhalation epochs were administered at hourly intervals, temperature was most consistently reduced in the combination condition and only declined in the THC (12.5 mg/mL) condition after the third exposure (**Figure 5**). Nicotine inhalation increased locomotor activity only in the nicotine-only condition in these two experiments and did not counter the locomotor suppressing effects of THC, where present in the 30-minute, 25 mg/mL experiment. The lowest exposure to THC, i.e. in the first 15 minute 12.5 mg/mL inhalation epoch, elevated activity compared with the vehicle inhalation but with further inhalation in the same session, THC suppressed activity. There was no evidence in that experiment that nicotine co-administration further increased the locomotor stimulation produced by the lowest dose of THC and no evidence nicotine could prevent the locomotor suppression associated with additional THC inhalation within the session.

As one minor caveat, while the present study did not seek to address tolerance there was an apparent reduction in sensitivity to the hypothermic effects of identical nicotine inhalation conditions from Experiment 1 to Experiment 3. Despite this, there was still hyperactivity observed in Experiment 4 in the nicotine-only condition and an apparently additive effect of nicotine and THC inhalation on the hypothermic response. The nicotine was active however additional investigation would be necessary to more precisely delineate any plasticity (tolerance or sensitization) to the effects of inhaled nicotine under various dosing paradigms.

In conclusion, these studies validate the use of this model for the study of the effects of ENDS based inhalation of nicotine in rats. We further show that combined inhalation of nicotine and THC produces effects that are most consistent with an interpretation of independent, rather than interactive, effects.

## Contributors

MAT and MJP designed the studies. MC created the vapor inhalation equipment that was used. MJP, TMK and ELH performed the research and conducted initial data analysis. MJP and MAT conducted statistical analysis of data, created figures and wrote the paper. All authors approved of the submitted version of the manuscript.

